# Sodium-glucose co-transporter 2 Inhibitors Act Independently of *SGLT2* to Confer Benefit for Heart Failure with Reduced Ejection Fraction in Mice

**DOI:** 10.1101/2024.04.29.591665

**Authors:** Justin H. Berger, Timothy R. Matsuura, Caitlyn E. Bowman, Renee Taing, Jiten Patel, Ling Lai, Teresa C. Leone, Jeffrey D. Reagan, Saptarsi M. Haldar, Zoltan Arany, Daniel P. Kelly

## Abstract

Sodium-glucose co-transporter 2 inhibitors (SGLT2i) are novel, potent heart failure medications with an unknown mechanism of action. We sought to determine if the beneficial actions of SGLT2i in heart failure were on- or off-target, and related to metabolic reprogramming, including increased lipolysis and ketogenesis. The phenotype of mice treated with empagliflozin and genetically engineered mice constitutively lacking SGLT2 mirrored metabolic changes seen in human clinical trials (including reduced blood glucose, increased ketogenesis, and profound glucosuria). In a mouse heart failure model, SGLT2i treatment, but not generalized SGLT2 knockout, resulted in improved systolic function and reduced pathologic cardiac remodeling. SGLT2i treatment of the SGLT2 knockout mice sustained the cardiac benefits, demonstrating an off-target role for these drugs. This benefit is independent of metabolic changes, including ketosis. The mechanism of action and target of SGLT2i in HF remain elusive.

---

Sodium-glucose co-transporter 2 inhibitors (SGLT2i), originally designed as drugs to treat diabetes, have profound benefit in treating heart failure (HF). Multiple cardiovascular outcome studies have demonstrated remarkable reduction in HF hospitalization rates and death in patients with HF with reduced ejection fraction (HFrEF).^1^ Despite rapid adoption of SGLT2i in American and European standard-of-care guidelines for all classes of HF, the mechanism of their benefit is unknown. By preventing glucose re-absorption in the kidney, SGLT2i confer improved insulin sensitivity, modest weight loss, and slight reduction in systolic blood pressure. However, the observed benefits of SGLT2i in HF are much greater than can be explained by these modest, pleiotropic effects. Recent studies suggest an off-target effect, with multiple plausible mechanisms remaining, including SGLT2i-induced ketogenesis and re-programming cardiac substrate utilization.^2,3^ In this study we sought to determine whether SGLT2i exerts therapeutic benefit via on- or off-target effects in a mouse model of HFrEF, and address whether ketosis contributes to the beneficial action in HF.

A range of SGLT2i doses and routes of administration have been used in rodent studies to date with variable efficacy. We conducted dose-ranging studies in C57BL/6NJ male mice to define empagliflozin (empa) pharmacokinetic exposure that approximates efficacious doses used in human clinical trials.^4^ When admixed in standard chow diet (LabDiet, #5001), a dose of 50 mg/kg/day, based on average intake of 4-5 grams of food per day per mouse, resulted in a peak plasma concentration equivalent to published human data (Fig. 1A). Within three days of therapy initiation, this dosing strategy results in a trend toward reduction in serum glucose, increased fasting ketosis, and profound glucosuria, without significant weight loss.

**Figure 1.**
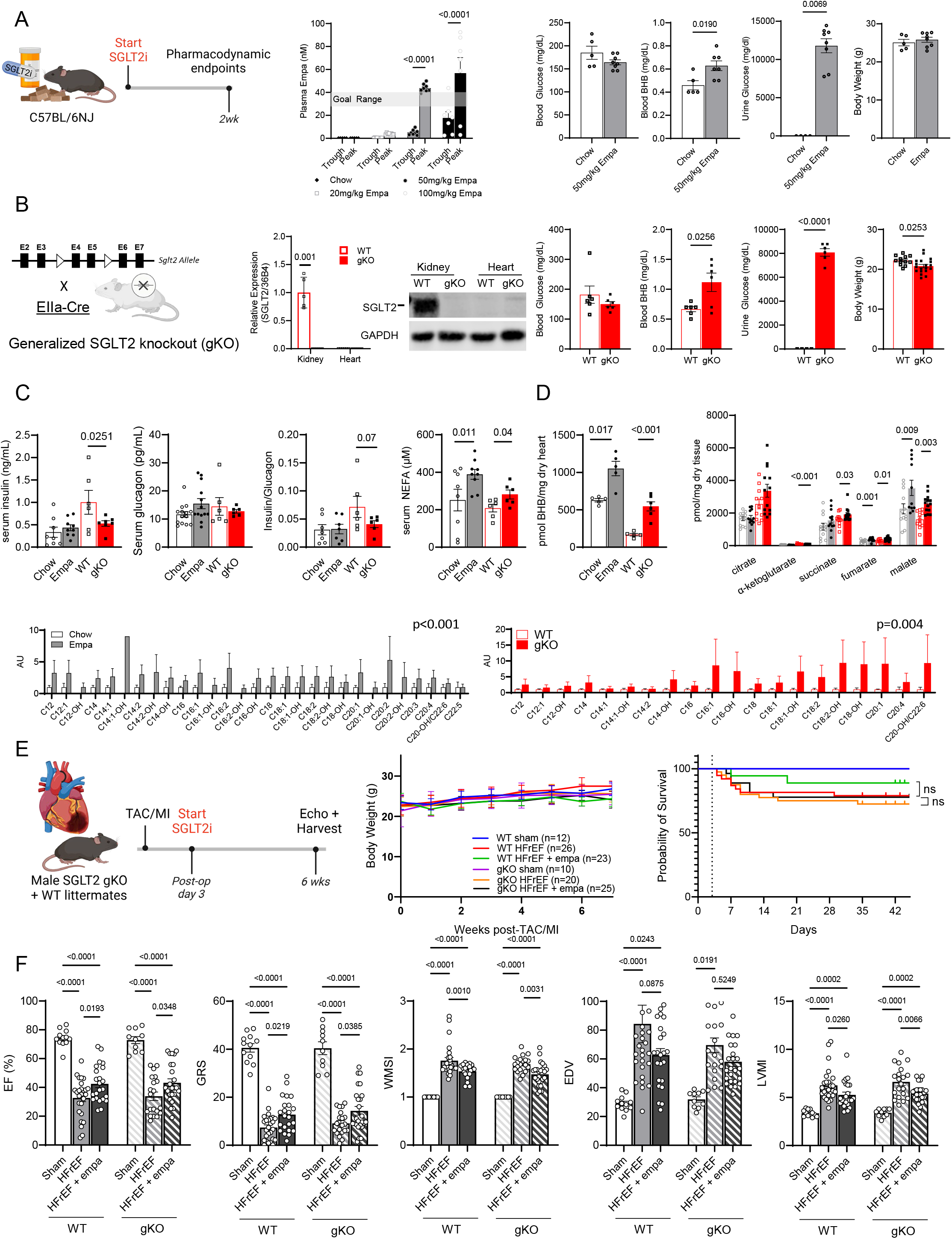
Pharmacologic and genetic SGLT2 inhibition have similar metabolic but divergent cardiac protective effects in HFrEF in mice. (A, left panel) Peak and trough plasma concentrations of empa following dose-ranging of admixed drug into chow diet of WT mice for 2 weeks (n=5-8 per group). (A, right graphs) Fasted blood glucose and beta-hydroxybutyrate (BHB) levels, urine glucose, and body weight (n=5-8 per group) in WT mice treated with 50 mg/kg/day empa for 2 weeks. (B, left panel) Generalized SGLT2 KO mouse with QT-PCR and representative western blot showing reduction in kidney and heart. (B, right graphs) Fasted blood glucose and BHB levels, urine glucose (n=6 per group), and body weight in WT and gKO littermates at age 8-10 weeks (n=12-18 per group). (C) Serum insulin, glucagon, insulin/glucagon ratio, and non-esterified free fatty acids, and (D) cardiac BHB concentration, targeted cardiac TCA cycle intermediates and cardiac acylcarnitine profile from 6-7 hour fasted WT +/- empa and WT vs gKO littermates (n= 8-14 per group). (E) Schema for TAC/MI +/- SGLT2i in WT and gKO littermates. (E, right) Weight trend and Kaplan-Meier curve, and (F) echo and gross morphologic endpoints for sham and TAC/MI +/- empa WT and gKO male littermates (n=10-12 sham, 20-26 HF per group). All data are presented as mean ± SEM and analyzed by Student’s two-tailed t-test (A-D) or 2-way ANOVA/Tukey multiple comparisons test (D – ACs and F) using Prism 9. P values shown. Abbreviations: BHB, beta-hydroxybutyrate; BiV/TL, biventricular weight to tibia length ratio; EDV, end-diastolic volume; ESV, end-systolic volume; EF, ejection fraction; empa, empagliflozin; GLS, global longitudinal strain; GRS, global radial strain; LVMI, left ventricular mass index (to pre-surgical weight); SGLT2, sodium-glucose co-transporter 2; SGLT2i, SGLT2 inhibitor; TAC/MI, transaortic constriction with apical myocardial infarction; TCA, tricarboxylic acid; WMSI, wall motion score index; WT, wildtype. Schema in A, B and E created with BioRender.com.

We sought to compare the metabolic and cardiac phenotype of SGLT2i-treated mice with mice lacking SGLT2. Generalized SGLT2 knockout (gKO) mice were generated by breeding SGLT2 floxed mice (MMRRC #049746-UCD) with mice ubiquitously expressing EIIa-Cre recombinase (JAX #003724) (Fig. 1B). gKO mice were born in expected Mendelian ratios, developed normally, and had no baseline cardiac phenotype (data not shown), though they were slightly smaller compared to WT littermates (Fig. 1B). The effects of empa treatment and of global SGLT2 deficiency on circulating metabolic parameters were similar, including a trend to mildly decreased fasting serum glucose, elevated fasting ketosis, and profound glucosuria compared to controls (Fig. 1B). We hypothesized that the benefit of SGLT2i via reduced insulin/glucagon ratios leads to changes in cardiac fuel delivery, including increased lipolysis and subsequent ketogenesis. Insulin was modestly reduced in the gKO but not empa-treated serum, while glucagon was unchanged and insulin/glucagon ratios trended lower with gKO (Fig. 1C). Circulating non-esterified fatty acids (NEFA) were increased in both empa-treated and gKO groups compared to matched controls, consistent with increased lipolysis. Targeted metabolomic analysis of left ventricle (LV) myocardium from gKO and empa-treated WT mice was also performed. Myocardial levels of the ketone body beta-hydroxybutyrate were increased approximately two-fold in both empa-treated and gKO conditions, consistent with the observed ketonemia (Fig. 1D). A subset of myocardial TCA cycle intermediates (malate, fumarate) was increased in a similar pattern with empa treatment and gKO. Lastly, medium- and long-chain acylcarnitines in the LV were significantly increased (2-way ANOVA, p<0.004) with empa treatment and gKO consistent with increased fatty acid utilization or a pathway “bottlenecking” effect. Taken together, these results indicate that SGLT2 KO largely phenocopies the metabolic effects of SGLT2i treatment.

To compare the impact of genetic SGLT2 deficiency and pharmacologic SGLT2 inhibition in the context of HFrEF, 8–10-week-old male SGLT2 WT and gKO littermates on a C57BL/6NJ background were subjected to trans-aortic constriction with apical myocardial infarction (TAC/MI), an established rodent HFrEF model,^5^ with and without empa treatment started on post-operative day 3 (Fig. 1E). Empa treatment and genotype did not affect body weight or survival. Following TAC/MI, WT mice treated with empa demonstrated improved LV ejection fraction (EF) and reduced pathological remodeling at six weeks (Fig. 1F). In contrast, gKO mice subjected to TAC/MI had no difference in EF, strain, or pathologic remodeling compared to WT controls. Moreover, treatment of the SGLT2 gKO mice with empa produced the same degree of benefit to cardiac functional and remodeling endpoints observed in treated WT controls.

In conclusion, generalized SGLT2 deficiency phenocopies key metabolic effects of pharmacological SGLT2i treatment with empa in mice including increased ketosis, lipolysis, and evidence of myocardial substrate utilization shifts. Despite similar metabolic profiles, global genetic SGLT2 deficiency did not recapitulate cardiac protection of empa therapy in an established HFrEF model. Beneficial effects of SGLT2i treatment in mice genetically lacking SGLT2 conclusively demonstrate that SGLT2 is not required for the *in vivo* mechanism of cardiac protection by SGLT2i in a HFrEF model. This study expands on recent work to demonstrate that in a physiologically relevant pre-clinical model of HFrEF, SGLT2i can exert therapeutic benefits via off-target pharmacology.^3^ The lack of benefit seen in gKO HFrEF mice, despite the presence of robust ketosis observed in this setting, argues against myocardial ketone body oxidation as a likely therapeutic mechanism. Further work remains to identify specific cardiac or extra-cardiac targets of SGLT2i that confer beneficial effects in HF.

## Acknowledgments

We thank Fang Xie and Brooke Rock at Amgen for technical assistance. The cardiac surgeries and echocardiography were performed by the Rodent Cardiovascular Phenotyping Core (RRID: SCR_022419) at the University of Pennsylvania supported by the Penn Cardiovascular Institute and NIH S10OD016393. Metabolomics studies were performed by the Metabolomics Core (RRID:SCR_022381) in the Penn Cardiovascular Institute and supported, in part, by NCI P30 CA016520.

